# Nationwide citizen science reveals core and rare osmotolerant yeast communities in Swedish honey

**DOI:** 10.64898/2026.10.21.760674

**Authors:** Karl Persson, Michael Martindale, Linda Winnicki, Agneta Persson, Cecilia Geijer

## Abstract

Honey represents an extreme microbial habitat, comprising >80% sugar, low pH, oxidative stress, and antimicrobial compounds. Despite these conditions, specialized yeast species persist in nectar and honey microbial communities, where their metabolic activity may contribute to honey maturation and chemistry. To characterize the diversity and functional traits of honey-associated yeasts, we conducted a nationwide citizen science survey with the help of Swedish beekeepers, yielding ∼2,400 isolates from 119 honey samples across 103 locations. Sequencing of ITS and LSU barcode regions of cultured isolates identified 85 species across 26 genera, dominated by *Entelexis*, *Starmerella*, *Zygosaccharomyces*, *Rhodotorula*, *Aureobasidium*, and *Metschnikowia*. While a small number of core taxa were consistently detected across geographic and environmental gradients, most species were rare and locally restricted, indicating a highly uneven community structure. Phenotypic assays of representative strains, typically one per isolated species, revealed broad osmotolerance, with several *Entelexis, Starmerella* and *Zygosaccharomyces* strains capable of growth at up to 80% glucose. Sugar utilization experiments showed widespread fructophilic metabolism, particularly under high-sugar conditions, and this trait was enriched among the most frequently recovered taxa. Selected isolates also fermented diluted honey, altering both its volatile profile and chemical composition. Together, these results demonstrate that honey hosts a taxonomically diverse but functionally specialized yeast community shaped by strong osmotic selection. This study establishes the largest culture-based collection of honey-associated yeasts to date, underscores the value of citizen science for large-scale microbial ecology, and identifies honey as an important reservoir of osmotolerant and fructophilic yeasts with ecological and biotechnological relevance.

## Introduction

Honeybees commonly forage within a few kilometers of their hive, although foraging distances vary considerably with season, landscape, and resource availability and can extend substantially farther [1]. Through these foraging trips, the bees continuously encounter and spread microorganisms among flowers, nectar, and the hive, where honey is produced and stored. Honey can therefore be viewed as a microbial reservoir receiving continuous inputs of microorganisms through bee activity. However, the high sugar concentration, low pH, oxidative environment, and antimicrobial compounds characteristic of honey create strong physiological filters that favor specialized, stress-tolerant microorganisms. Thus, despite these harsh conditions, honey and the hive environment support diverse microbial communities composed of bacteria and fungi, including both commensal organisms and pathogens. For example, lactic acid bacteria are common in fresh honey and the bee gut, where they contribute to fermentation and inhibit competing microbes [2]. Both unicellular yeasts and filamentous fungi are frequently introduced via pollen and nectar and may influence honey maturation and quality [3–5]. The hive can also host bee-pathogens such as *Paenibacillus larvae*, the agent of American foulbrood [6], and fungi causing chalkbrood [7]. Collectively, these microorganisms form a microbial network that shapes honey composition and hive ecology.

Given that yeasts are common inhabitants of the hive environment and honey, identifying which taxa persist in this extreme habitat is important for several reasons. Ecologically, honey-associated yeasts can influence plant–pollinator interactions and hive processes by shaping microbial communities during nectar concentration and honey maturation, with consequences for bee nutrition and health [8, 9]. From a food perspective, honey has long been studied as a reservoir of spoilage yeasts, particularly osmotolerant genera such as *Zygosaccharomyces*, which can affect the stability of sugar-rich products [10]. Other recurrent genera, including *Starmerella* and *Metschnikowia*, are increasingly recognized for their roles in nectar and honey fermentation and interactions with pollinators [8, 11]. Finally, honey represents an underexplored source of biotechnologically relevant yeasts, as many honey-associated genera exhibit high osmotolerance and metabolic traits producing valuable compounds under high-sugar conditions [12]. Together, these ecological, food-related, and biotechnological perspectives provide complementary motivations for characterizing the yeast communities that inhabit honey.

Although several honey yeasts have been reported, as summarized in reviews such as [13], we still lack a comprehensive understanding of their diversity across landscapes and environments. Large-scale surveys are limited, geographic coverage remains sparse, and many potentially novel yeast species remain undescribed. Large-scale yeast bioprospecting has demonstrated the potential to uncover previously uncharacterized species and strains with valuable physiological traits [14]. Moreover, the functional traits of honey yeasts, such as their ability to grow at extreme sugar concentrations, produce valuable metabolites, and influence bee health, are only superficially understood [12, 13, 15]. To address these gaps, we initiated a nationwide citizen science project engaging Swedish beekeepers to collect honey samples for yeast isolation. Sweden provides a suitable system for large-scale sampling of honey-associated yeasts, combining a well-established beekeeping network with pronounced environmental gradients from southern agricultural regions to northern boreal landscapes, enabling the collection of geographically and ecologically diverse samples. Our objective was to establish a large and diverse collection of honey-derived yeasts, and to characterize their taxonomic and ecological diversity and evaluate phenotypic traits relevant to stress tolerance and honey fermentation.

In this study, we report on the isolation of ∼2,400 yeast strains from 119 honey samples collected across Sweden. Using ITS and LSU barcode sequencing of cultured isolates, combined with growth characterization and analytical chromatography, we characterized the taxonomic diversity of honey-associated yeasts and identified strains with remarkable osmotolerance and fermentation capacity. This work establishes the largest nationwide collection of honey-associated yeasts to date, providing new insights into their ecological diversity and adaptation to high-sugar environments.

## Results

### Nationwide citizen-science sampling reveals a large culturable yeast collection

To explore the diversity of yeasts associated with honey, we leveraged a citizen science approach, engaging beekeepers across Sweden to participate in the collection of honey samples (Fig. 1). Beekeepers were recruited through outreach on social media platforms, where a recruitment post shared through our own networks was subsequently circulated within beekeeping communities. The post provided a brief overview of the research goals and invited beekeepers to contribute honey samples to the study (Fig. 1A). Within a few days of the announcement, more than 130 beekeepers expressed interest in participating. To standardize sample collection, participants received sampling kits containing sterile, barcoded tubes, instructions, metadata questionnaires, and prepaid return envelopes (Fig. 1B).

**Figure 1.**
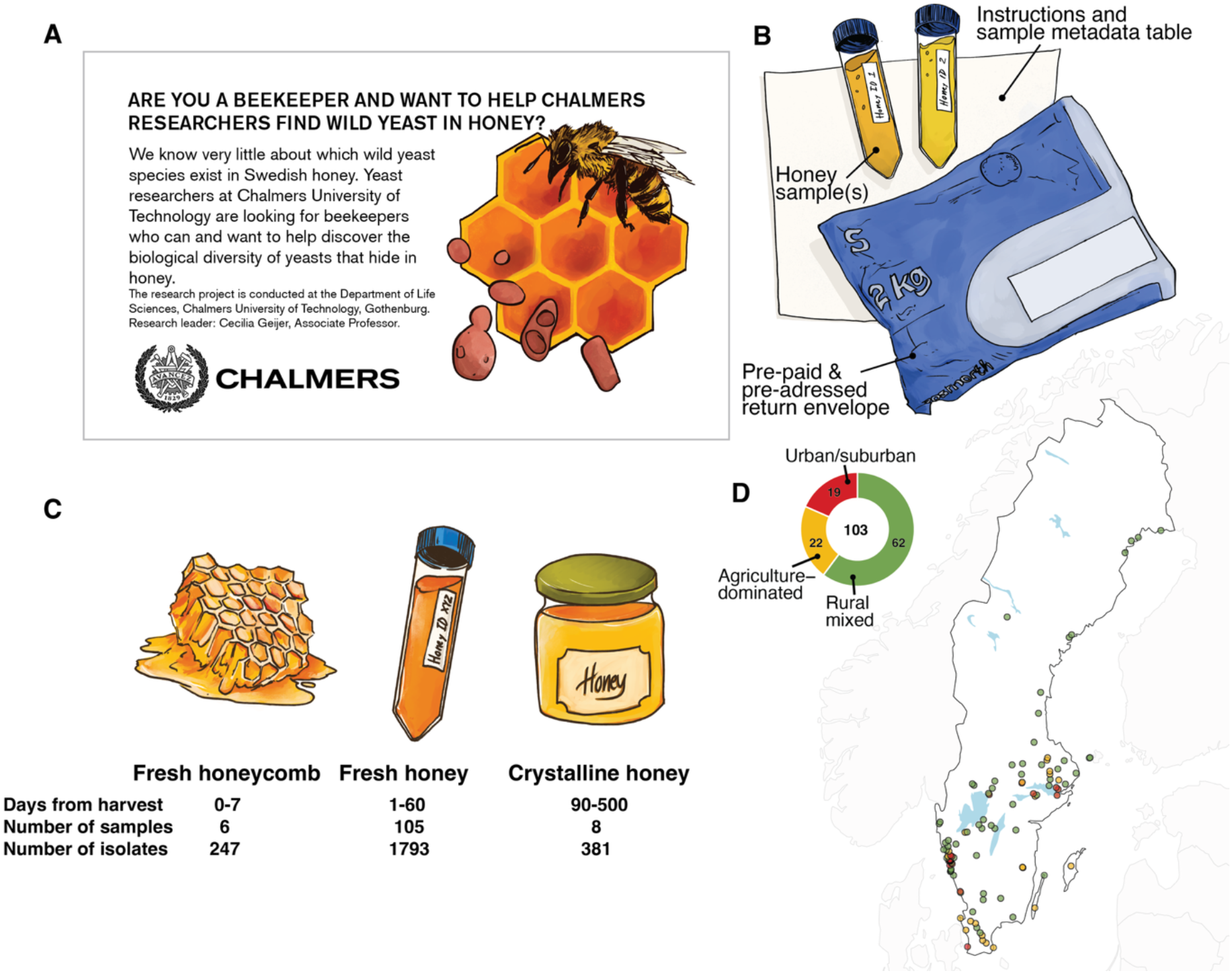
Citizen science–based sampling of honey across Sweden. A) Advertisement used to recruit participating beekeepers (translated from Swedish). B) Returned sampling kit as received from participants, including completed metadata forms and barcoded tubes filled with honey samples. C) Types of honey samples collected, with sample counts indicated. D) Map describing the geographic distribution of the 103 sampling locations. Points are color-coded according to three major environments, with proportions summarized in an inset donut chart.

Most of the participating beekeepers collected honey from one site (geographic location), while 15 beekeepers sampled honey from two sites that were located at least 2 km from each other (median distance of 6.4 km) and one beekeeper provided samples from 5 unique locations within the Gothenburg area. Additionally, from one sampling site, we collected 12 samples over a longer time span (3 years) and seasonal span (4 months). Out of 119 samples, 105 were freshly extracted honey, eight were crystalline honey, and six were freshly cut honeycombs (Fig. 1C). Most samples came from Buckfast bees (77), with some reported as crosses with local bee types. Other bee types included Italian honeybees (*Apis mellifera ligustica*, 19), Carniolan honeybees (*A. m. carnica*, 9), and Nordic honeybees (*A. m. mellifera*, 1). Bee type was unspecified for 13 samples. Upon receipt, each sample was catalogued using the barcoded identifiers, and metadata were digitized for integration with the sample dataset. Overall, this initiative successfully gathered honey from a wide geographic range, enabling a comprehensive study of 2421 culturable and isolated yeast strains from 103 unique geographic locations (Fig. 1D). The sites were grouped into three broad environment classes (urban/suburban, agriculture-dominated and rural mixed), where the majority represented mixed landscapes, and forests were present to various degrees around all sampled hive sites.

### Honey hosts a taxonomically diverse and uneven yeast community

Following method optimization, yeasts were isolated using concentration plating on antibiotic-supplemented medium (Fig. 2A; Supplementary Methods). Yeast-like colonies were purified and identified by ITS sequencing, supplemented with LSU sequencing when necessary. The isolation approach identified 85 unique species belonging to 26 genera, with the majority (83%) of the isolates belonging to the Ascomycota phylum. The strain collection is dominated by yeast species from the genera *Entelexis* (39%), *Zygosaccharomyces* (18%), *Starmerella* (9%), *Rhodotorula* (9%), *Aureobasidium* (7%), *Metschnikowia* (6%), *Filobasidium* (3%) and *Naganishia* (3%) (Fig. 2B), all of which have been reported previously from nectar, honey, or other bee-associated products [4, 13, 22, 23]. The taxonomic tree in Figure 2C illustrates the taxonomic relationships of the identified species and highlights the diversity of the honey yeast collection. While most isolates were true yeasts (2119/2421, 87.4%), the total collection was more morphologically diverse, with some taxa classified as yeast-like, dimorphic fungi or filamentous fungi.

**Figure 2.**
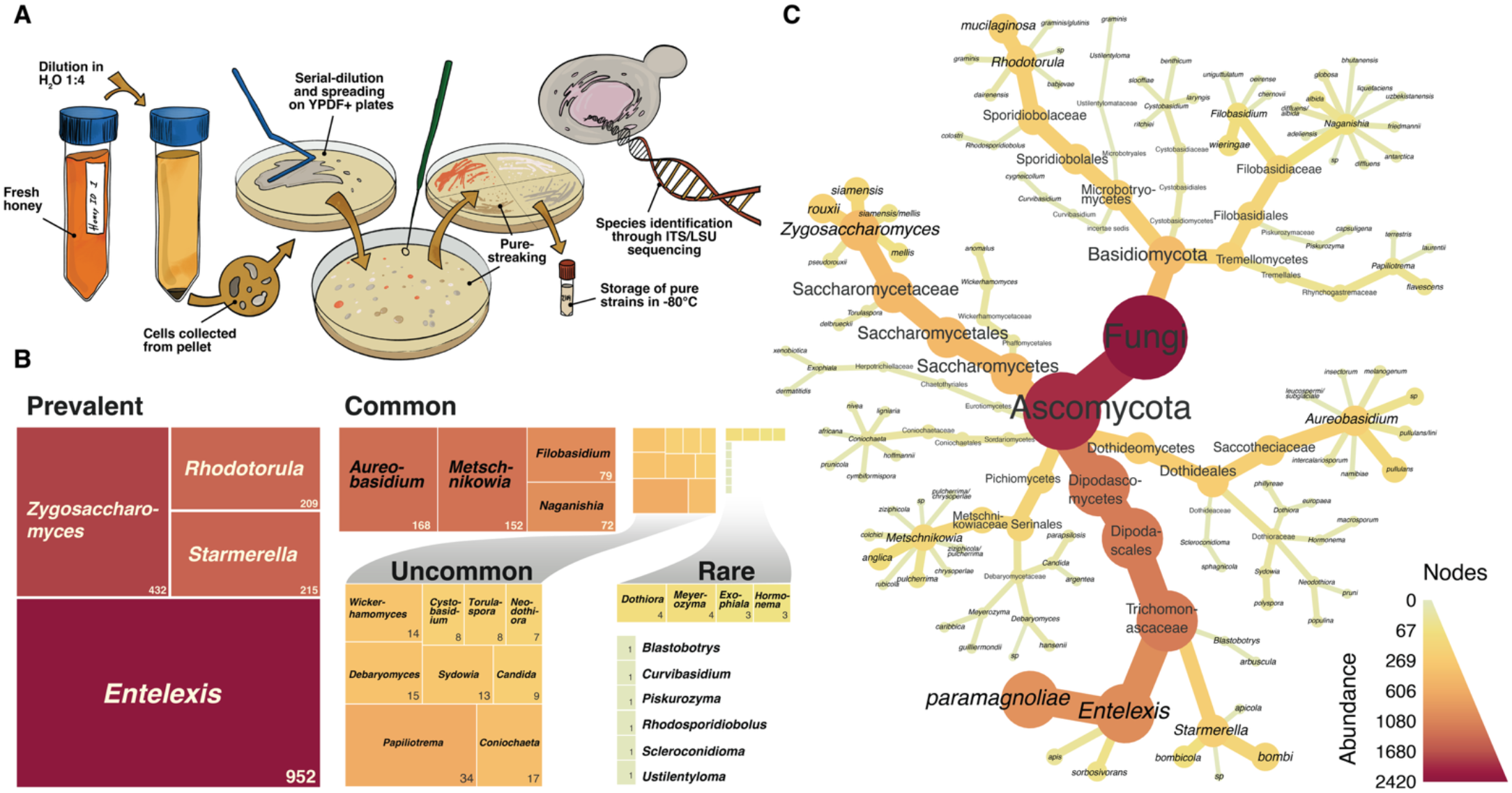
Taxonomic identification and diversity of honey-derived yeasts. A) Workflow for yeast isolation, including dilution of honey, centrifugation-based concentration of cells, plating on selective medium, colony purification, DNA sequencing, and strain preservation. B) Treemap of yeast genera with rectangle area proportional to relative abundance; uncommon and rare genera are grouped and scaled for clarity. C) Taxonomic overview of identified strains across hierarchical levels (phylum to species). The layout is not scaled to evolutionary distance. Branch lengths reflect relative diversity, whereas node size and color intensity are proportional to the abundance of each taxon.

A summary of species prevalence and diversity is provided in Table S1. The most abundant yeast in our collection was *Entelexis paramagnoliae*, which represented 37.9% of all isolates (Fig. 2 C, Table S1). Notably, *Entelexis* was only recently separated from the genus *Starmerella* following taxonomic revision [24], and *E. paramagnoliae* was previously classified as *Starmerella magnoliae* and, even earlier, *Candida magnoliae*. This species has previously been reported from honey and nectar environments and is known for its strong osmotolerance, consistent with adaptation to high-sugar substrates [12, 13]. Several additional *Entelexis* species (*E. apis* and *E. sorbosivorans*) and relatively closely related *Starmerella* species (i.e. *S. bombi* and *S. bombicola*) were also detected, further highlighting the prominence of these genera within the honey-associated yeast community. The second most abundant species was *Zygosaccharomyces rouxii* (8.3% of isolates), a common honey-associated osmotolerant yeast [12], followed by *Rhodotorula mucilaginosa* (8.0%), which is widespread on plants and in nectar [25]. *Metschnikowia* species were also recurrent, consistent with their frequent association with flowers and pollinating insects [11, 13]. In contrast, pathogenic yeasts including several *Candida* and *Cryptococcus* species commonly associated with human hosts (e.g., *C. albicans*, *C. auris*, *C. neoformans*) were largely absent from our collection, suggesting that the extreme osmotic conditions of honey impose strong environmental filtering on yeast communities. Moreover, we did not isolate any *Saccharomyces cerevisiae* strains, despite recent reports of this species from honey [23]. This may reflect regional climatic differences between studies, as temperature is an important determinant of *Saccharomyces* species distributions [26].

We also recovered several rarely reported taxa, including *Piskurozyma capsuligena* and *Curvibasidium cygneicollum*, for which we found no previous association with honey. A subset of isolates also showed less than 95% similarity to reference sequences in the database. Although some low-identity strains can be attributed to short and low-quality sequence data, others likely represent novel taxa to be characterized in a follow-up study.

### Core taxa recur across geographic regions while many other species were infrequently detected

Sweden spans more than 1,500 km from south to north, encompassing temperate to subarctic climates and a wide diversity of floral resources. Agricultural areas are often dominated by mass-flowering crops, whereas forested regions provide nectar and pollen from wildflowers and flowering trees. Floral resources also vary seasonally, from early-flowering *Salix* spp. to fruit trees, berry-producing plants, and late-season heather (*Calluna vulgaris*). Most honey samples (90%) were collected during summer (29 in June, 49 in July, and 24 in August) from diverse urban, agricultural, and forested landscapes (Fig. 1D).

Geographic mapping of yeast sample origins revealed that the distribution of yeast genera varied across locations and sample types, as illustrated by the relative genera abundance within each sample (Fig. 3A). Two locations were sampled more intensively to capture additional yeast diversity: a rural site (Site 1, Fig. 3A; 502 isolates, 25 species) and an urban site near the Gothenburg Botanical Garden (Site 2, Fig. 3A; 160 isolates, 24 species). Sampling density was higher in southern regions of Sweden, consistent with a greater representation of active beekeepers in these areas. The most abundant genera (including *Entelexis*, *Starmerella*, *Rhodotorula* and *Zygosaccharomyces*) were consistently found across multiple sites, suggesting that these yeasts are key components in the honey microbiome (Fig. 3B). Similarly, the three most common species, *E. paramagnoliae*, *R. mucilaginosa* and *Z. rouxii*, were found in multiple samples, occurring in 82, 39 and 39 samples, respectively (Table S1). In contrast, a large fraction of the species detected in our dataset were rare, with 53 of 85 species represented by five or fewer isolates, which were grouped in the “other” category in Figure 3A. Although most of these taxa were confined to single sampling locations, they still contribute to the overall ecological diversity of the honey-associated mycobiota.

**Figure 3.**
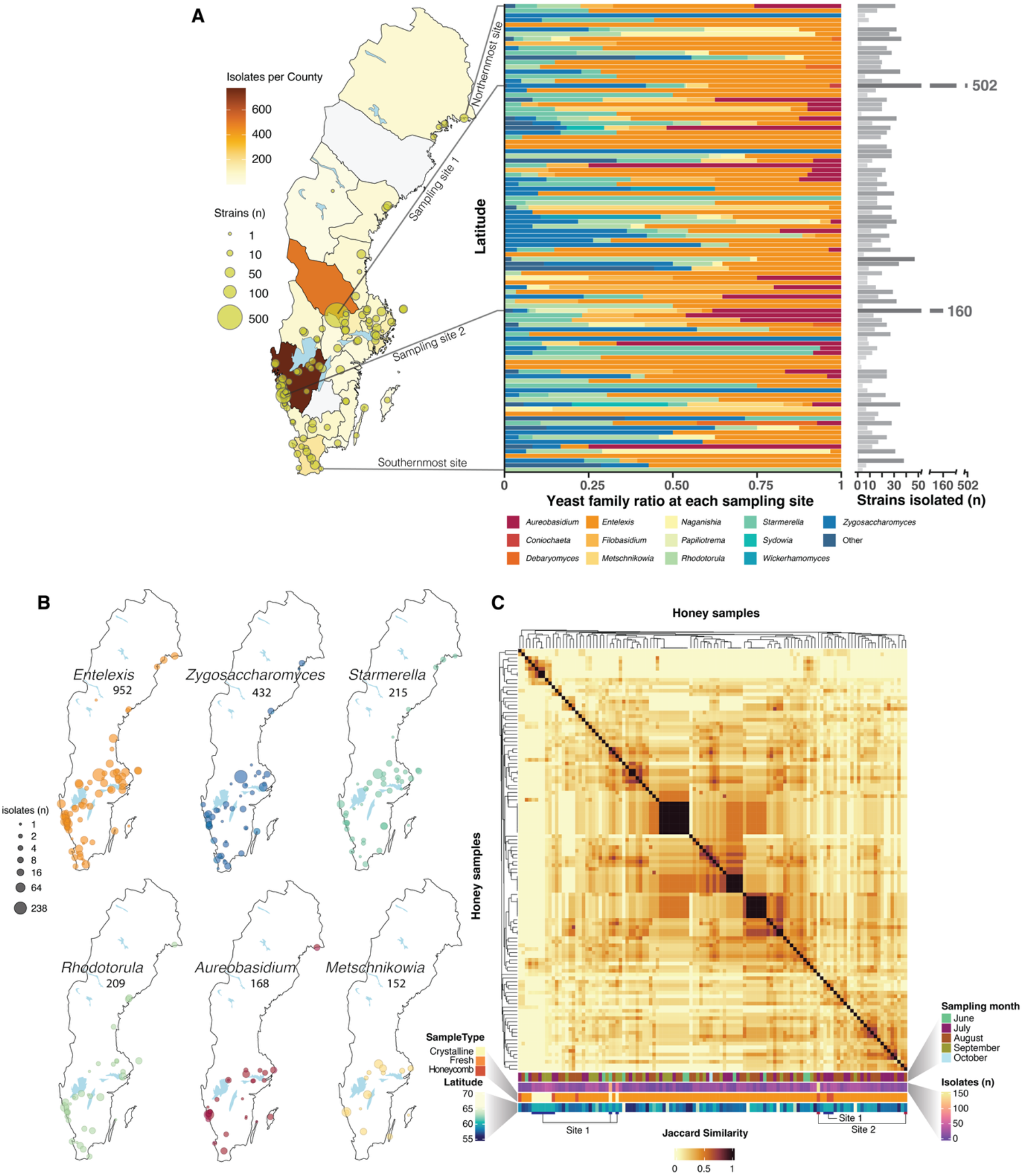
Diversity and geographic distribution of honey-associated yeasts across Sweden. A) Overview of sampling sites and taxonomic composition. The map shows the geographic distribution of all sampling locations. Bar plot shows the relative abundance of the 12 most abundant yeast genera across sites and the remaining grouped into “other”, ordered by latitude. The number of isolates obtained per site is shown in grayscale to the right. B) Geographic distribution of the six most abundant yeast genera, showing the sampling sites where each genus was detected. Species are indicated by the same colors as in A. C) Clustered heatmap of pairwise Jaccard similarity among honey batches based on yeast species presence–absence. Rows and columns were ordered by hierarchical clustering of the Jaccard distance matrix. Annotations indicate sampling month, sample type, latitude, number of isolates per batch, and site identity for the two intensively sampled locations (Site 1 and Site 2).

We compared yeast species composition among honey batches using Jaccard similarity coefficients visualized as a clustered heatmap (Fig. 3C). Overall, similarity was low to moderate, indicating substantial heterogeneity among samples, with only a few localized clusters of similar communities. These clusters showed no consistent association with sampling month or latitude. Some grouping by sample type was observed, particularly for honeycomb and crystalline honey samples, although this should be interpreted cautiously due to uneven sampling. Together, the results suggest that honey-associated yeast communities are shaped by multiple factors.

### Honey yeasts exhibit broad tolerance to extreme sugar concentrations

A representative panel of 89 strains was selected for phenotypic characterization, typically one per species, with additional isolates from abundant taxa, including *E. paramagnoliae* (n = 8) and *Metschnikowia pulcherrima* (n = 5). These strains originated from 49 sampling sites across Sweden (Fig. 4A). As a first step, we assessed growth on the main honey sugars glucose and fructose. Most species (87.3%) grew in 2% glucose and fructose (Fig. 4B), while some grew only at higher sugar concentrations and others failed to grow detectably in liquid culture despite growing on agar. These differences likely reflect variation in osmotic tolerance, growth kinetics, and assay conditions. Given that high sugar concentrations impose strong osmotic stress on the microbiome in honey, we next examined growth across a gradient of glucose concentrations (Figure 4B). In media containing glucose at increasing concentrations from 10% to 80% (w/v), most isolates grew robustly at 10% and 20%, whereas higher concentrations (40%–80%) markedly reduced growth (Figure 4B). The most osmotolerant yeasts were various *Entelexis*, *Starmerella* and *Zygosaccharomyces* strains, which sustained growth up to 80%. *Zygosaccharomyces* strains generally displayed shorter lag phases than the *Entelexis* and *Starmerella* strains at glucose concentrations of 60-80% (mean 53.0 h vs. 80.6 h), which is consistent with previous findings [5]. *Debaryomyces hansenii* and several of the *Metchnikowia* strains were able to grow at glucose concentrations of up to 70%. In contrast, genera such as *Cystobasidium*, *Dothiora*, and *Naganishia* were among the most sensitive to elevated glucose concentrations. Overall, these results highlight substantial interspecies variation in osmotolerance, with only a subset of taxa adapted to withstand the extreme sugar concentrations of 70-80%.

**Figure 4.**
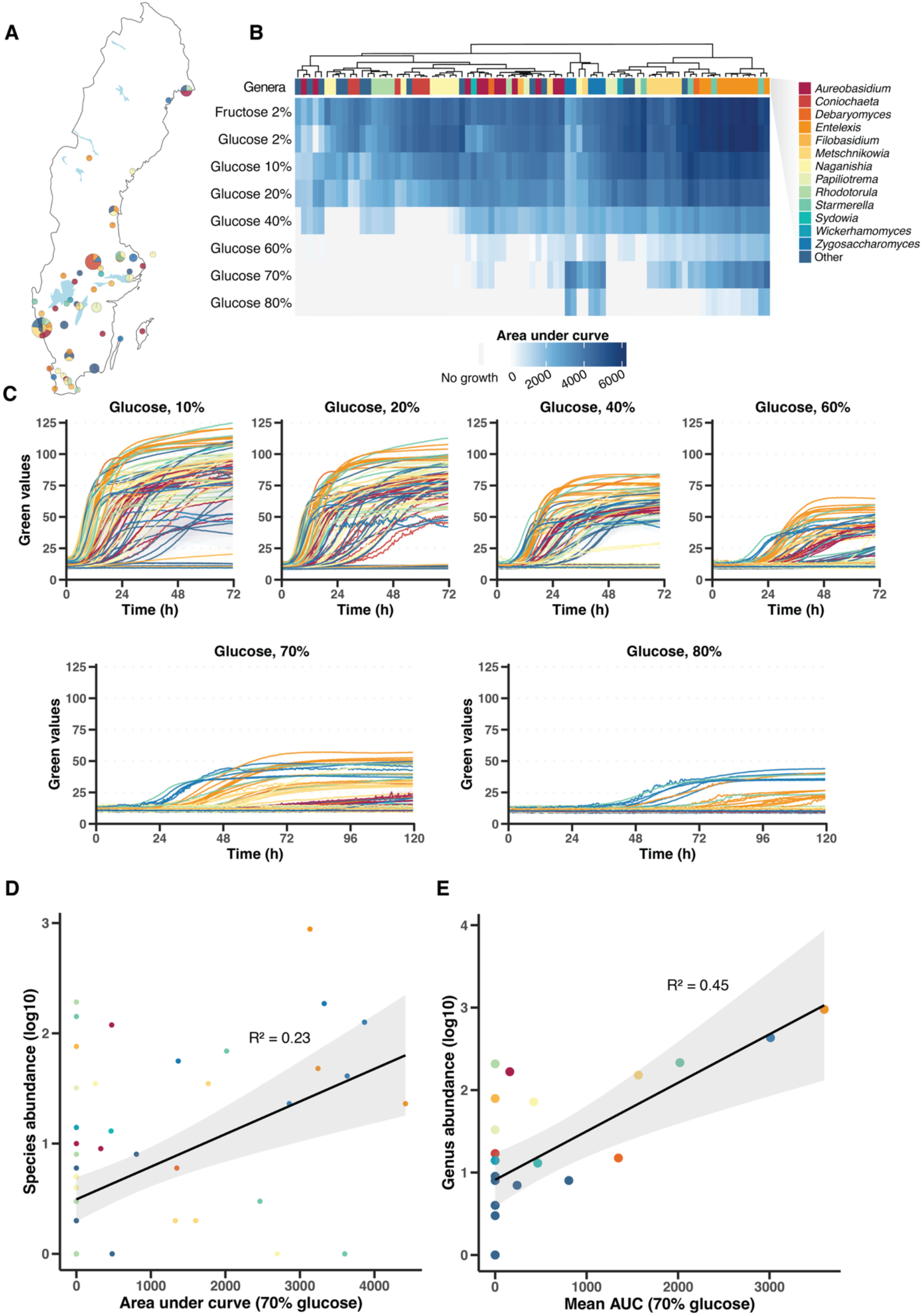
Growth characteristics of the core yeast collection. A) Geographic distribution of the core collection. Pie charts represent sampling sites, with slices corresponding to individual strains colored by genus. B) Heatmap showing growth performance across conditions, expressed as area under the curve (AUC). Rows represent growth conditions and columns represent strains. C) Mean growth curves of strains across a gradient of glucose concentrations (10–80%). Lines are colored by genus. Growth was monitored in 96-well microtiter plates using a Growth Profiler, with Green Values derived from image analysis used as a proxy for biomass accumulation. D) Relationship between species abundance and growth performance at 70% glucose, measured as area under the growth curve (AUC). The black line represents the linear regression fit, with the gray shaded area indicating the 95% confidence interval; R^2^ denotes the coefficient of determination. E) Same as in D, but with abundance aggregated at the genus level. Points/curves in (A–E) are colored according to genus, as indicated in B.

We hypothesized that the dominance of specific yeast taxa in honey is associated with traits that facilitate growth under increasing osmotic stress during nectar concentration and honey maturation, such that more abundant taxa would exhibit greater osmotolerance. To test this, we examined the relationship between isolate abundance and growth performance across increasing glucose concentrations. While this relationship was variable at the strain level (Fig. 4D), a clear pattern emerged when data were aggregated at the genus level (Fig. 4E). *Entelexis*, *Starmerella* and *Zygosaccharomyces* were all highly abundant and capable of sustained growth at high sugar concentrations, *Metschnikowia* occupied an intermediate position, and less frequently represented genera clustered at lower abundance and lower tolerance to osmotic stress. Together, these results demonstrate that capacity to grow in high concentrations of sugar contributes to the prevalence of specific yeast lineages in honey, consistent with increasing osmotic stress acting as an environmental filter during honey maturation.

### Fructophilic metabolism is enriched among dominant honey yeasts

Most yeasts preferentially consume glucose when exposed to both glucose and fructose simultaneously [27]. However, honey is composed of high concentrations of both fructose and glucose, with fructose concentrations typically exceeding glucose [28, 29]. This environment may favor yeasts capable of efficiently utilizing fructose in the presence of glucose, either through co-utilization of both sugars or through preferential fructose consumption. Notably, a fructophilic phenotype, defined by preferential utilization of fructose, has been reported in several yeasts from the genera *Starmerella* and *Zygosaccharomyces* [13, 30, 31], both of which are well represented in our collection. To assess sugar preferences across the honey yeast collection, we cultured 89 representative strains in mixed-sugar media containing either 2% glucose + 2% fructose or 10% glucose + 10% fructose. The lower concentration serves as a baseline under relatively mild osmotic conditions, whereas the higher concentration more closely reflects nectar and maturing honey.

In the 2% + 2% condition, most strains consumed glucose more rapidly than fructose over the course of the experiment (46 h) (Fig. 5A, B). However, several strains, primarily within *Zygosaccharomyces*, *Starmerella* and *Entelexis* (e.g., *Z. rouxii*, *Z. pseudorouxii*, *Z. siamensis*, *S. bombi*, *E. paramagnoliae* and *E. apis*), exhibited clear fructophilic phenotypes. Under the 10% + 10% condition, strains that were fructophilic at 2% + 2% displayed an even stronger preference for fructose at higher concentrations (Fig. 5A, C). Additionally, several strains from other genera (eg., *Aureobasidium*, *Coniochaeta*, *Dothiora*, *Filobasidium*) showed approximately equal consumption of glucose and fructose, and some strains (e.g., *Blastobotrys*, *Rhodotorula*, *Naganishia*) even exhibited weak fructophilic tendencies (Fig. 5C). Interestingly, we note a substantial overlap between strains capable of growth in 70-80% glucose and those exhibiting a fructophilic phenotype (i.e *Zygosaccharomyces*, *Starmerella* and *Entelexis*) (Fig. 4C and S1). Together, these findings indicate that sugar utilization patterns in honey-associated yeasts are concentration dependent and suggest a positive association between fructophilic behavior, extreme sugar tolerance, and species prevalence in honey.

**Figure 5.**
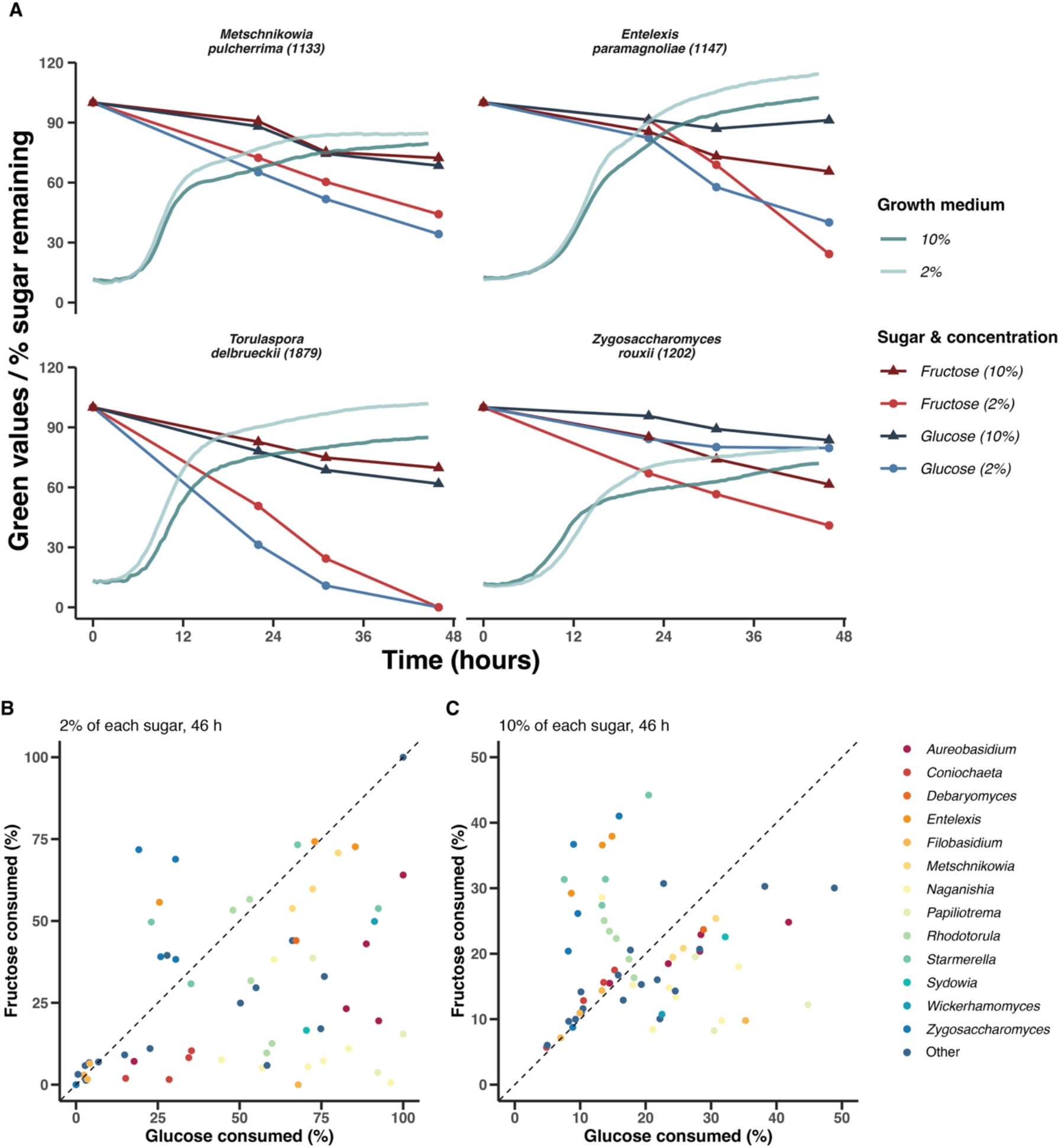
Sugar utilization patterns of honey-associated yeasts. A) Time-resolved growth and HPLC measurements of glucose and fructose consumption in representative yeast strains (complete dataset shown in Fig. S2). Growth was monitored using Green Values, while glucose and fructose concentrations are expressed as percentages of their initial concentrations over 46 h. Each strain was cultivated in multiple parallel wells, with separate wells sampled at each time point for sugar quantification. B) Relationship between glucose and fructose consumption in media containing 2% glucose + 2% fructose after 46 h. Each point represents one strain; points above the diagonal indicate preferential fructose utilization, whereas points below indicate preferential glucose utilization. C) Relationship between glucose and fructose consumption in media containing 10% glucose + 10% fructose after 46 h. Points in (B– C) are colored by genus.

### Osmotolerant yeasts can ferment diluted honey

The ability of many isolates to grow at exceptionally high sugar concentrations raises the possibility that honey-associated yeasts are not merely passive contaminants but may actively shape honey chemistry. To assess their capacity to alter the honey composition under conditions conducive to growth and metabolic activity, we selected ten strains representing abundant and highly osmotolerant *Zygosaccharomyces, Starmerella*, *Entelexis* and *Metschnikowia* species from our collection. The strains were evaluated in a model fermentation system consisting of honey diluted to 60%, 70%, 80%, or 90%. All strains produced detectable gas during the 25-day incubation period in up to 80% honey, and several strains also did so at 90% honey, indicating active fermentation. However, both the onset and extent of fermentation varied among strains and honey concentrations (Fig. 6A).

**Figure 6.**
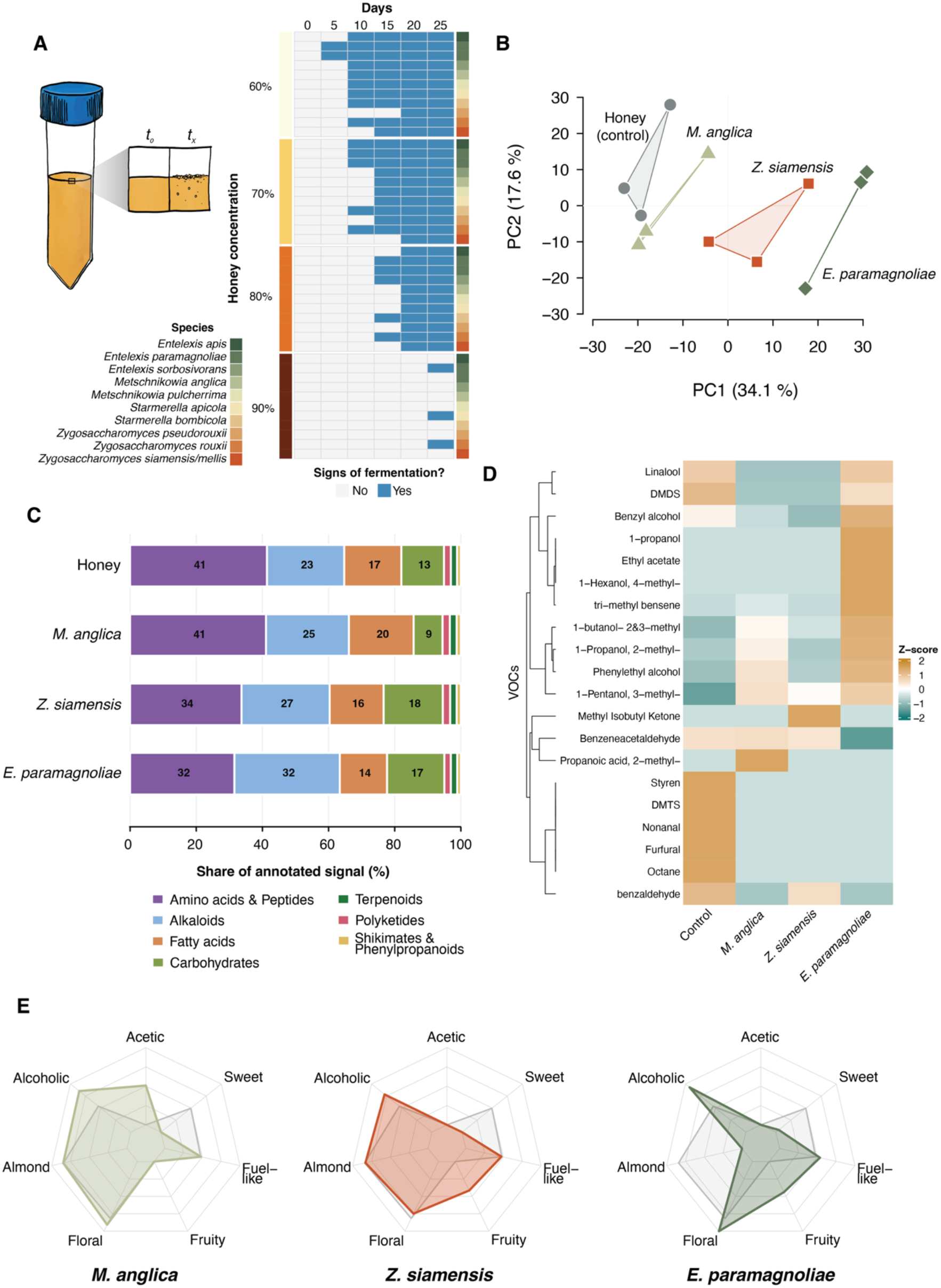
A model fermentation experiment demonstrating metabolic potential of honey-associated yeasts. A) Apparent fermentation of honey by selected osmotolerant yeast strains in diluted honey (60-90% (v/v), scored every 5 days over 25 days; blue, fermentation observed. B) PCA of the fermented-honey metabolome (LC-MS/MS, positive mode; 1571 curated features, log2-transformed and Pareto-scaled). Polygons denote three biological replicates. C) CANOPUS (NPClassifier) pathway composition of the same features, as each pathway’s share of total annotated signal (714 features with pathway probability > 0.5). D) Heatmap of VOCs detected from fermented honeys (GC-MS) as per-compound z-scores, rows hierarchically clustered. E) Radar plots showing log10-transformed summed odor activity values (OAVs) of the VOCs shown in (D), illustrating the relative contribution of different aroma-associated compound groups to the predicted scent profile.

To investigate whether yeasts produce distinct extracellular metabolite profiles, we performed untargeted HPLC-MS/MS analysis of fermented honey after 21 days for three different yeast species, namely *Z. siamensis*, *E. paramagnoliae*, and *M. anglica*. Among the three strains, *E. paramagnoliae* showed the highest apparent fermentation activity at the time of sampling, based on visual inspection, while *Z. siamensis* and *M. anglica* displayed moderate fermentative performance (data not shown). Thus, the results should be viewed as snapshots of strain-specific fermentation trajectories, as the different yeasts had acted on the diluted honey to varying degrees by the time of sampling. Of 3,589 aligned features retained after blank and intensity filtering, 1,571 passed quality control (QC-RSD ≤ 0.4, D-ratio ≤ 0.6) and were carried forward. PCA analysis revealed clear separation of the individually fermented honeys from each other and from the non-fermented honey control (Fig. 6B), with treatment accounting for 53 % of the total variance (PERMANOVA *p* = 0.0002). Classification of annotated metabolites into pathways revealed that fermentation altered the relative distribution of metabolite categories, although the annotated signal remained dominated by amino acids and peptides, alkaloids, carbohydrates and fatty acids in all samples. Amino acids and peptides accounted for 41% of the annotated signal in the uninoculated honey but declined to 34% and 32% after fermentation with *Z. siamensis* and *E. paramagnoliae*, respectively, while remaining unchanged in *M. anglica*-fermented honey (41%). Comparatively, alkaloids showed a reciprocal pattern, rising from 23% of the annotated signal in the uninoculated honey to 27% and 32% in *Z. siamensis* and *E. paramagnoliae*-fermented honey, respectively, while fatty acid share varied comparatively little. Relative to the uninoculated control (13%), carbohydrates decreased in *M. anglica* (9%) but increased in *Z. siamensis* (18%) and *E. paramagnoliae* (17%). Although the specific sensory consequences remain to be determined, changes in the abundance and composition of metabolite classes such as amino acids, fatty acids, and alkaloids could potentially influence flavor development and contribute to strain-specific sensory properties of fermented honey [32–34].

To assess the impact of fermentation on aroma, the volatilomes of 70% diluted honey fermented for 21 days by the three yeasts were analyzed by GC–MS. Fermentation resulted in pronounced shifts in both the presence and abundance of volatile organic compounds (VOCs) for all yeast-inoculated honeys relative to uninoculated controls, indicating strong yeast-driven modulation of the honey volatilome (Fig. 6D). Higher alcohols constituted the dominant class of VOCs and increased markedly following yeast fermentation, with the most pronounced effect observed for *E. paramagnoliae.* Aromatic compounds such as phenylethyl alcohol were also strongly enriched, whereas several aldehydes present in the control remained relatively stable or even decreased. In contrast, esters and ketones exhibited more species-specific production patterns.

Odor activity value (OAV) analysis, based on compound abundance relative to odor detection thresholds, revealed that a smaller subset of compounds, including higher alcohols and aromatic aldehydes, contributed substantially to the predicted aroma profiles (Fig. 6E, Table S2). These profiles differed markedly among the tested yeasts, suggesting species-specific contributions to aroma development. Fermentation by *M. anglica* was associated with enrichment of several higher alcohols and aromatic compounds linked to acetic and alcoholic sensory characteristics (Fig. 6E).

*Z. siamensis* produced a comparatively distinct but less pronounced VOC profile, characterized by a broader distribution of alcohols and aldehyde-associated compounds, corresponding to moderate fruity and alcoholic sensory attributes. In contrast, *E. paramagnoliae* fermentation was associated with pronounced floral, alcoholic and fruity aroma characteristics. Together, these findings indicate that different honey-associated yeasts can drive distinct aroma trajectories during honey fermentation.

## Discussion

Honey supported a surprisingly diverse yeast community despite its extreme osmotic conditions. Our nationwide citizen science survey yielded ∼2,400 honey-associated yeast isolates representing 85 species across 26 genera, providing broad geographic coverage and a highly diverse strain collection. Consistent with previous studies [15, 35–37], *Entelexis*, *Zygosaccharomyces*, and *Starmerella* were the dominant genera, followed by *Rhodotorula*, *Aureobasidium*, and *Metschnikowia*. Their widespread occurrence across geographically distinct regions suggests that honey imposes strong ecological filters that favor a specialized community of osmotolerant yeasts. While a small number of taxa were common across many samples, most species were rare and locally restricted, including several isolates with low sequence similarity to known species that may represent undescribed taxa. This pattern of a few dominant taxa coexisting with many rare species resembles that observed in other microbial ecosystems [38, 39], and highlights honey as a reservoir of underexplored fungal diversity.

Growth assays revealed that most honey yeasts grow in glucose concentrations of ≥60% (w/v), consistent with strong physiological constraints associated with high osmolarity in honey. Such extreme osmotolerance is uncommon among yeasts outside specialized high-sugar and high-salt habitats [40], but has been repeatedly documented in honey-associated genera such as *Zygosaccharomyces* and *Starmerella* [15]. The ability of these taxa to persist under high osmolarity conditions likely explains their dominance across geographically distant honey microbiomes. Interestingly, several isolates from genera not typically associated with honey (i.e *Aureobasidium*, *Filobasidium*) also displayed substantial osmotolerance, suggesting that the capacity for growth under high-sugar conditions extends across a broader range of yeast lineages than previously appreciated. Most striking, however, was the overwhelming dominance of *E. paramagnoliae*, which alone accounted for more than one third of all isolates. This pattern aligns with its frequent detection in honey and nectar worldwide [31, 41], but the scale and geographic breadth of our sampling further emphasizes its strong ecological association with honey.

Beyond osmotolerance, the isolates exhibited diverse metabolic strategies. While many strains preferentially consumed glucose under low-sugar conditions, several species including *Z. rouxii*, *Z. siamensis*, *S. bombi*, and *E. paramagnoliae* displayed clear fructophilic phenotypes. Fructophily is well documented in *Starmerella* and *Zygosaccharomyces*, where a preference for fructose over glucose is thought to confer an advantage in nectar and honey [13, 42]. Interestingly, we found that strains from several additional genera showed approximately equal consumption of glucose and fructose, and weak fructophilic tendencies. These findings suggest that reduced glucose preference, or even fructophily, may be more phylogenetically widespread than previously recognized, highlighting the potential for convergent evolution of sugar-utilization strategies in high-sugar environments. Moreover, fermentation experiments showed that *Z. siamensis*, *E. paramagnoliae*, and *M. anglica* differ markedly in their ability to transform diluted honey, generating distinct volatile and non-volatile metabolite profiles. Because these profiles represent only a snapshot of a dynamic process, future studies incorporating temporal sampling will be needed to resolve the ecological significance of yeast-mediated transformations during nectar ripening and honey maturation. The pronounced species-specific differences observed here highlight both the limited understanding of secondary metabolism in honey-associated yeasts and their potential as sources of novel bioactive compounds and honey-derived fermented products, including specialty meads, vinegars, and fermented foods [43].

While our study provides a unique culture-based view of honey-associated yeasts, it also has limitations. Culture-dependent approaches inherently exclude non-culturable taxa and likely underestimate overall diversity. Existing metagenomic studies on honeybee-associated microbiomes has focused primarily on bacteria and viruses [44], leaving fungal diversity comparatively underexplored. Integrating metagenomic or amplicon-based approaches would enable a more comprehensive view of the yeast community, including uncultured and low-abundance taxa, although distinguishing active honey-associated yeasts from environmental DNA remains challenging. In addition, our sampling was largely restricted to a single season, limiting insights into temporal dynamics. Seasonal variation in floral resources, nectar composition, and bee foraging behavior may influence the structure of honey-associated yeast communities. Future studies combining high-throughput culturing, metagenomics, and functional genomics will be essential to resolve the full diversity and ecological roles of honey-associated yeasts.

Taken together, our results indicate that honey acts as a powerful ecological filter that selects for a specialized yeast community dominated by a few highly adapted taxa but also harbors rare and potentially novel lineages, highlighting its role as a unique reservoir of microbial diversity. The distinct physiological and metabolic capabilities of different honey-associated yeasts suggest that they can contribute to honey maturation processes while also representing promising resources for biotechnological applications. By linking large-scale community patterns with functional traits, this study provides a foundation for future work exploring how honey-associated yeasts influence honey chemistry, hive ecology, and bee-flower-microbe interactions.

## Methods

### Citizen science sampling of honey

Beekeepers across Sweden were recruited through social media outreach on Facebook and LinkedIn, where our recruitment post was shared within beekeeping communities. Approximately 130 beekeepers expressed interest, of whom the first 100 were selected to participate. Participants received standardized sampling kits containing sterile, barcoded 50 mL collection tubes, sampling instructions, and questionnaires documenting hive location, honey collection date, bee type, and surrounding environmental conditions. Beekeepers contributed not only honey samples and environmental metadata but also observations and suggestions through ongoing communication with the research team. Collected samples were returned by post, registered using their unique barcodes, and stored at room temperature until processing.

### Yeast Isolation from Honey Samples

Following an initial comparison of direct plating, liquid enrichment, and concentration plating (Supplementary Methods), the latter was selected for yeast isolation because it provided higher recovery of culturable yeasts while avoiding potential enrichment bias. Honey samples (8–14 g) were diluted threefold (w/w) in sterile distilled water, homogenized, and centrifuged at 5,000 × g for 10 min to concentrate microbial cells. The supernatant was removed, and the resulting pellet was resuspended in the remaining liquid. Two consecutive tenfold dilutions were prepared, and 100 µL aliquots were spread onto YPDF+ agar plates. The medium contained 1% yeast extract, 2% peptone, 2% glucose, 2% fructose, and 2% agar (w/v), supplemented with chloramphenicol and ampicillin (100 mg/L each) to suppress bacterial growth. Plates were incubated at approximately 25 °C for 3–7 days, and colonies displaying yeast-like morphology were selected and purified by repeated streaking on fresh YPDF agar. Purified isolates were subsequently grown in liquid YPDF medium and cryopreserved in 25% (v/v) glycerol at −80 °C.

### Species identification and strain selection

Genomic DNA was extracted from purified yeast isolates using an alkaline lysis method. The internal transcribed spacer (ITS) region was amplified by PCR using primers ITS1 (5’-TCCGTAGGTGAACCTGCGG-3’) and ITS4 (5’-TCCTCCGCTTATTGATATGC-3’). For isolates where ITS amplification was unsuccessful, the large subunit (LSU) rRNA region was amplified using primers NL-1 (5’-GCATATCAATAAGCGGAGGAAAAG-3’) and NL-4 (5’-GGTCCGTGTTTCAAGACGG-3’). PCR products were purified and Sanger sequenced by Eurofins Genomics using ITS4 and NL-4 as sequencing primers for the respective regions. Species identification was performed using BLAST searches against the NCBI ITS, LSU, and standard nucleotide databases. For each sequence, the top 3–5 matches were compared based on sequence identity, query coverage, bit score, and E-value. When database annotations were ambiguous or uninformative, additional high-scoring matches were examined to determine the most likely identification. Species assignments were guided by a minimum sequence identity threshold of 98.41% to reference sequences [16]. For isolates falling below this threshold, the species name of the closest matching reference sequence was provisionally assigned, and sequence identity values were retained to facilitate future taxonomic investigation. Sequences of insufficient quality or containing ambiguous base calls were excluded from further analysis. A representative panel of 89 strains was selected for phenotypic characterization, typically one per species, with additional strains included for selected abundant taxa to capture potential intraspecific variation. Selection also considered morphological diversity and geographic origin. Detailed molecular protocols are provided in Supplementary Methods.

### Growth characterization

Growth of the 89 representative yeast strains was characterized in 96-well microtiter plates using a Growth Profiler 960 (Enzyscreen). Strains were precultured in YPDF medium at 25 °C for 4 days, harvested by centrifugation, and washed with YP medium (1% yeast extract, 2% peptone). Washed cells were diluted tenfold in YP medium, and 10 µL of the diluted suspension was inoculated into microtiter wells containing 240 µL of YP medium supplemented with either 2% fructose or glucose at concentrations of 2%, 10%, 20%, 40%, 60%, 70%, and 80% (w/v). To investigate sugar utilization preferences, strains were additionally cultivated in mixed-sugar media containing either 2% glucose + 2% fructose or 10% glucose + 10% fructose (w/v), with samples collected for subsequent HPLC analysis. Cultures were prepared in quadruplicate and incubated at 30 °C with shaking at 250 rpm for up to 7 days. Growth was monitored by automated imaging, with Green Values derived from image analysis used as a proxy for biomass accumulation. Growth curves were analyzed to determine parameters including lag phase duration, maximum growth rate, final yield, and area under the growth curve (AUC). Detailed cultivation procedures are provided in Supplementary Methods.

### Fermentation of Honey

To assess the fermentation capacity of selected honey-associated yeasts, commercial honey (Svensk Honung, Svensk Honungsförädling AB) was heat-treated at 90 °C for 1 h to liquefy the crystallized honey and inactivate endogenous microorganisms, then cooled overnight at room temperature. The honey was diluted to 60%, 70%, 80%, and 90% (v/v) with sterile deionized water and thoroughly homogenized. Aliquots of 10 mL were dispensed into sterile 15 mL tubes. Ten yeast strains selected for their high osmotolerance were recovered from YPDF agar plates and inoculated in triplicate into each honey dilution using one loopful of cells (∼10 µL). Uninoculated heat-treated honey (negative control), a time-zero sample (T0), and actively fermenting honey (positive control) were included. Cultures were incubated in sealed tubes at 25 °C in darkness without agitation, and fermentation was monitored qualitatively by visual inspection for gas formation over 25 days. In a separate fermentation experiment, strains were cultivated in 70% honey under the same conditions. Samples were collected on day 21 for analysis of volatile organic compounds (GC–MS) and extracellular metabolites (UPLC–MS/MS). Additional experimental details are provided in Supplementary Methods.

### Chromatography and mass-spectrometry analyses

### HPLC analysis of extracellular metabolites

Sugar consumption was assessed in the mixed-sugar cultures described above by sampling separate culture wells at 0, 22, 31, and 46 h. Culture supernatants were collected by centrifugation, filtered through 0.2 µm filters, and analyzed using a JASCO HPLC system equipped with UV and refractive index detectors and a Rezex ROA Organic Acid H+ column. Glucose, fructose, and other extracellular metabolites were identified using analytical standards and quantified from calibration curves. Chromatograms were processed using ChromNAV 2.05 software. Additional analytical conditions are provided in Supplementary Methods.

### Untargeted LC–MS/MS analysis of fermented honey

Extracellular metabolites were extracted from fermented honey collected on day 21 using ethyl acetate following acidification to pH 3. Metabolite profiling was performed using an ACQUITY Premier UPLC system coupled to a Xevo MRT mass spectrometer (Waters). Mass spectra were acquired using electrospray ionization in positive and negative modes. Raw data were processed using MS-DIAL v4.6 for peak detection, alignment, and MS/MS deconvolution, followed by filtering, normalization, and annotation using a dedicated metabolomics pipeline. Features were filtered based on blank signals, detection frequency, and quality-control criteria, retaining 1,571 features for downstream analysis. Compound classes were predicted using SIRIUS/CANOPUS [17–19]. Detailed analytical conditions and processing parameters are provided in Supplementary Methods.

### GC–MS analysis of volatile compounds

Volatile organic compounds (VOCs) were analyzed from fermented honey collected on day 21 using headspace solid-phase microextraction (SPME) coupled to GC–MS. Samples were equilibrated at 40 °C for 10 min, followed by VOC extraction using a PDMS/DVB/CAR SPME fiber for 20 min at 40 °C. Chromatographic separation was performed using an Agilent GC–MS system equipped with a DB-WAX column. Compounds were identified based on mass spectra and retention times, and quantification was performed using flame ionization detection (FID). Odor activity values (OAVs) were estimated by relating compound abundances to published odor detection thresholds in air [20, 21]. Because SPME extracts only a fraction of the volatile compounds present in the headspace, these values were interpreted as relative estimates of odor activity rather than absolute sensory thresholds. Detailed analytical procedures and OAV calculations are provided in Supplementary Methods.

### Data analysis and visualization

Statistical analyses and data visualization were primarily performed in R (version 4.6.1). Growth performance was quantified using area under the growth curve (AUC), and relationships between yeast abundance and osmotolerance were examined using linear regression. Community similarity among honey samples was assessed using Jaccard similarity indices and hierarchical clustering. Metabolomic differences among fermentation treatments were evaluated using principal component analysis (PCA) and permutational multivariate analysis of variance (PERMANOVA). Data visualization was performed using packages from the tidyverse and additional specialized libraries. Scientific illustrations were created digitally by hand, and final figure layouts were assembled and refined using Adobe Illustrator. Detailed analytical procedures and software packages are provided in Supplementary Methods.

## Acknowledgements

This study was supported by Formas grants 2023-01931 and 2025-00747, Chalmers Area of Advance Energy, and the Adlerbertska Foundation.

We thank Susanne Nilsson for helping us with public outreach and communication, including improvements to our social media material and news articles related to the project. We thank Daniel Brunnsåker at Chalmers Mass Spectrometry Infrastructure (CMSI) for his help with the acquisition and processing of the LC–MS/MS data, and Patrik Libander at Research Institutes of Sweden (RISE) for his help with the acquisition and processing of the GC–MS data. We also thank Jonas Warringer and Rike Stelkens for their thoughtful comments and critical feedback during manuscript preparation.

Thanks also to all beekeepers for their generous donation of honey samples: Agneta & Gunnar Persson, Albin Wågnert, Alexandra Kivelä, Anders Svensson, Andreas Carlsson, Andreas Hennius, Andreas Niklasson, Andreas Rebert, Anna Forsberg, Anna Gund, Aviva Hansson, Barbro Martinsson, Birthe Löfqvist, Björn Albinsson, Christer Berggren, Christer Dahlin, Christoffer Svenske, Daniel Bodin, David Hagerberg, David Hanson, Ebba Örtengren, Emil Nyholm, Eva Eriksson, Giulia Ray, Gunnel Engström, Haidi Andersson, Heléne Kennedal, Honungs Janne, Hugo Flordal, Jennifer Axelsson, Joacim Kämpe, Jocum Wigerup, Johanna Kylin, Johanna Vargvik, Johannes Lundgren, Jonathan Othén, Judith van der Weiden, June Hedfors, Karin Carolusson, Karina Öhrberg, Karl-Johan Holmstrand, Kerstin Namuth, Klas Liden, Lena Friberg, Lena Wyrner, Lillemor Bulukin, Line Obermeier, Linnéa Sandman Kruus, Lis-Mari Öhlund, Lisa Hansson, Malena Lidar, Malin & Tobbe Husberg, Malin Hansson, Margareta Strömberg, Margareta Westin, Marie Nilsson, Marika Pettersson, Markus Johansson, Martin Östling, Mats Ullberg, Mikael & Birgitta Leymann, Mirjam & Robert Landholm, Nils Andersson, Nils Jansson, Peter Jensen, Peter Svanberg, Pär Mollberg, Rasmus Anduri, Rebecka Gunnarson, Robert Niemi, Sabina Brandin, Simon Nilsson, Stefan Jacobsson, Stig Andersson, Stina Karlsson, Susan de Freitas, Susanne Hultström, Therese Sichan-Jansson, Thomas Axelsson, Tina Tyrchan, Tommy Gullisryd, Ulf Berthag, Ulf Stenback, Åke Lindahl, Åsa & Göran Olsson, Åse Johansson

During the preparation of this manuscript, the authors used ChatGPT (OpenAI) and Microsoft Copilot to assist with language refinement, phrasing, coding, and manuscript editing. All scientific interpretations and conclusions were developed by the authors, who critically reviewed and edited the AI-assisted text and take full responsibility for the final manuscript.

## Data availability

The datasets and R scripts underlying the analyses and figures presented in this study are publicly available at https://github.com/karl-persson/HoneyYeasts. Additional strain information, including DNA barcode sequences, is provided in Supplementary sequence table.

