## Supplementary material for "Nationwide citizen science reveals core and rare osmotolerant yeast communities in Swedish honey": Figure S1

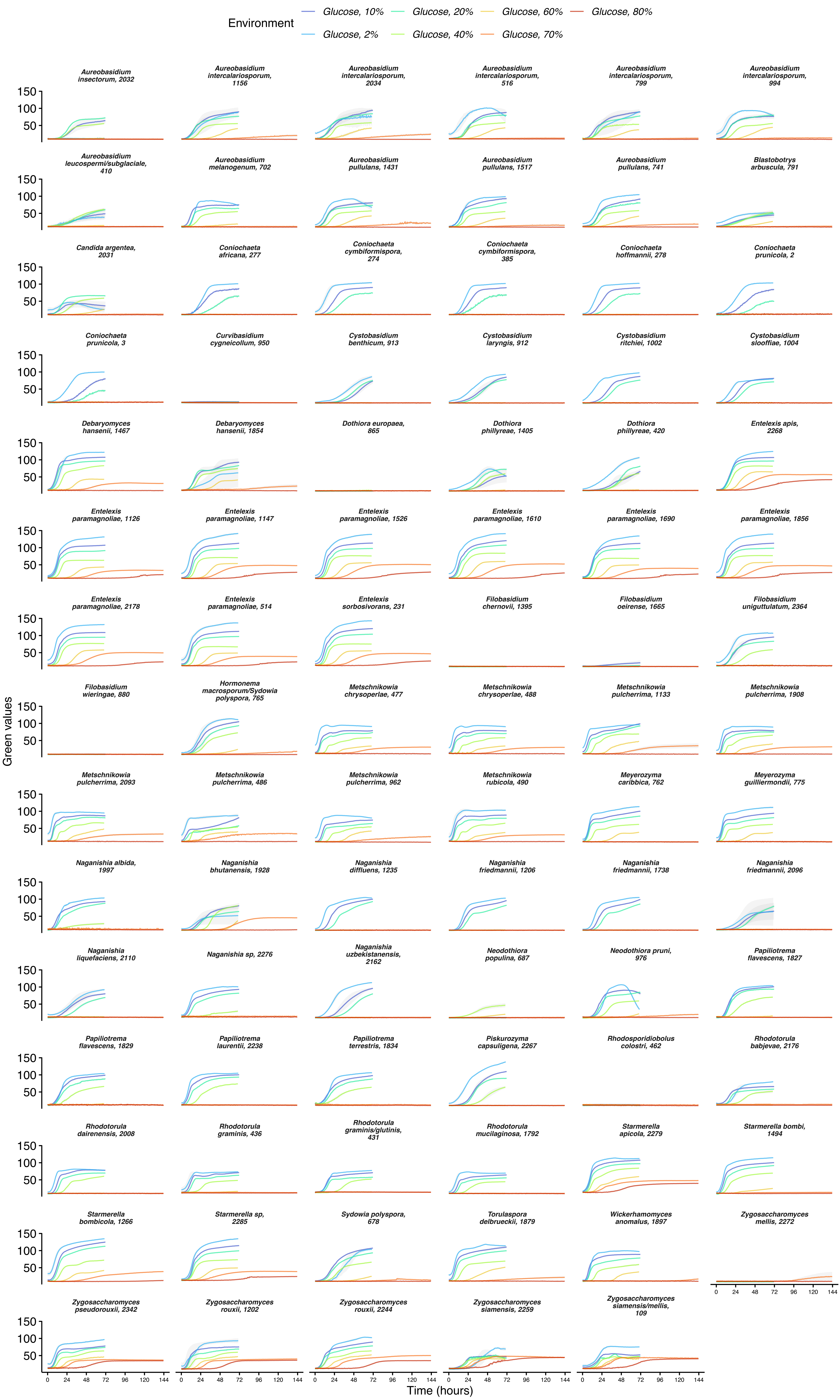


**Figure S1.** Growth curves of all strains across a gradient of glucose concentrations (2–80%). Each panel shows the growth the mean growth curve of individual strains (indicated by species and strain ID) under the 7 conditions. Growth curves were measured in 96-well microtiter plates using a Growth Profiler 960, which automatically images each well and quantifies changes in green pixel values as a proxy for biomass accumulation throughout cultivation. Curves represent three independent biological replicates per strain and condition.
