## Supplementary material for "Nationwide citizen science reveals core and rare osmotolerant yeast communities in Swedish honey": Figure S2

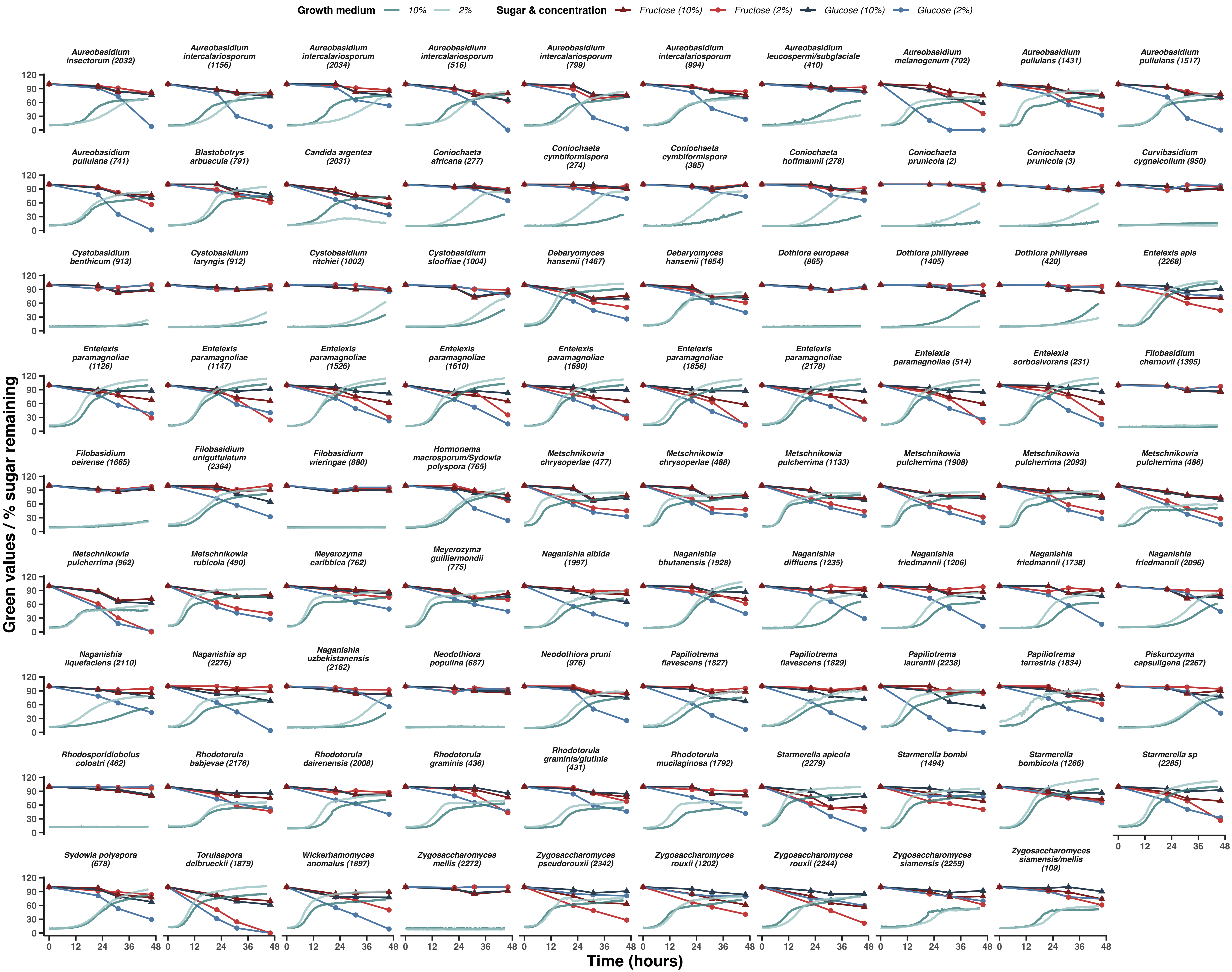


**Figure S2. Time-resolved growth and sugar utilization profiles of honey-associated yeasts.** Growth and glucose and fructose consumption were monitored over 46 h in mixed-sugar media containing either 2% glucose + 2% fructose or 10% glucose + 10% fructose (w/v). Each panel represents an individual yeast strain, identified by species name and strain ID. Growth was monitored using Green Values, while glucose and fructose concentrations are expressed as percentages of their initial concentrations. Lines distinguish growth and sugar consumption under the two experimental conditions.
