## Supplementary Methods for "Nationwide citizen science reveals core and rare osmotolerant yeast communities in Swedish honey"

### Optimization of yeast isolation

An initial pilot study was conducted using stored crystalline honey collected over several seasons from a single sampling location (Sampling site 1, fig 3A). Three approaches were evaluated for isolating culturable yeasts: direct plating, liquid enrichment, and concentration plating.

**Cultivation medium.** Yeasts were cultivated on a modified yeast extract–peptone–dextrose medium supplemented with fructose (YPDF), containing 1% yeast extract, 2% peptone, 2% glucose, 2% fructose, and 2% agar (w/v). To suppress bacterial growth, chloramphenicol and ampicillin were added at 100 mg/L each (YPDF+).

**Direct plating.** Honey was diluted tenfold in sterile distilled water, and 100 µL aliquots were spread onto YPDF+ agar plates. Plates were incubated at approximately 25 °C until colonies appeared.

**Liquid enrichment.** Honey was diluted 1:1 in liquid YPDF+ medium and incubated at approximately 25 °C for 24–72 h or until visible growth occurred. Cultures were subsequently diluted tenfold in sterile water and plated onto YPDF+ agar.

**Concentration plating.** Approximately 10 g of honey was mixed with three times its weight of sterile distilled water in 50 mL tubes and centrifuged at 5,000 × g for 10 min. The supernatant was discarded, and the pellet containing microbial cells and insoluble particles was resuspended in the remaining liquid. Two consecutive tenfold dilutions were prepared in sterile water and plated onto YPDF+ agar. Plates were incubated at approximately 25 °C for 3–7 days.

**Method selection.** Direct plating generally yielded few colonies, whereas liquid enrichment produced higher colony numbers but risked favoring rapidly growing species. Concentration plating provided consistently higher recovery without an enrichment step and was therefore selected for subsequent sample processing. Colonies were monitored to avoid overgrowth by filamentous fungi, and selected yeast-like colonies were purified by repeated streaking.

### Citizen science recruitment and honey sampling

**Recruitment of participants**. Swedish beekeepers were recruited through social media outreach on Facebook and LinkedIn. A recruitment post describing the research objectives and inviting participation was shared through the researchers' networks and subsequently circulated within beekeeping communities, including closed Facebook groups. Approximately 130 beekeepers expressed interest, of whom the first 100 were selected to participate.

In addition to collecting honey samples, participants provided information about their hives and surrounding environments and engaged in communication with the research team throughout the study. Several participants contributed observations and suggestions that informed sampling and the contextual interpretation of the collected data.

**Sampling kits and metadata collection.** Each participant received a standardized sampling kit containing two sterile, pre-barcoded 50 mL collection tubes, detailed sampling instructions, a metadata questionnaire, and a padded, prepaid return envelope.

The questionnaire collected information on hive location (geographic coordinates), honey extraction date, colony characteristics (including queen age and honeybee type), and the surrounding environment. Participants were instructed to collect approximately 45 mL of freshly extracted honey per tube, taking care to minimize contamination during sampling.

**Sample handling and storage.** Honey samples were sealed in the provided collection tubes and returned to the laboratory by post together with the completed questionnaires. Upon receipt, samples were registered using their unique barcodes, and the accompanying metadata were digitized and linked to the corresponding samples. Missing or unclear information was clarified through direct communication with participants when possible. Samples were stored at room temperature until processing.

### Yeast isolation and strain preservation

**Sample preparation.** Honey samples (8–14 g) were mixed with three times their weight of sterile distilled water and manually shaken until fully homogenized. Samples were adjusted to equal volumes with sterile distilled water and centrifuged at 5,000 × g for 10 min. The supernatant was carefully removed, and the resulting pellet was resuspended in the remaining liquid. For samples containing honeycomb fragments, floating wax was removed after centrifugation.

**Plating and incubation.** Two consecutive tenfold dilutions were prepared from the resuspended pellet using sterile distilled water. Aliquots (100 µL) were spread onto YPDF+ agar plates and incubated at approximately 25 °C for 3–7 days.

**Colony selection and purification.** Following incubation, colonies displaying yeast-like morphology were selected based on colony appearance and the absence of visible hyphal growth. Selected colonies were transferred to fresh YPDF agar without antibiotics and purified by at least two consecutive streaks to obtain pure cultures.

**Long-term strain preservation.** Purified isolates were inoculated into 3–5 mL YPDF broth and cultivated at approximately 25 °C with shaking at 150 rpm for 3–4 days. Cultures were supplemented with glycerol to a final concentration of 25% (v/v) and stored at −80 °C.

### DNA extraction, sequencing, and species identification

**Genomic DNA extraction.** Genomic DNA was extracted from purified yeast isolates by alkaline lysis. Briefly, 10 µL of sedimented cells from liquid cultures were suspended in 50 µL of 10 mM NaOH and heated at 99 °C for 20 min. Samples were vortexed briefly and centrifuged to pellet cellular debris. The resulting supernatant was used as the template for PCR (1 µL per reaction).

**PCR amplification.** PCR was performed in 25 µL reactions using Phusion High-Fidelity DNA Polymerase. The internal transcribed spacer (ITS) region was amplified using primers ITS1 (5'-TCCGTAGGTGAACCTGCGG-3') and ITS4 (5'-TCCTCCGCTTATTGATATGC-3'). For isolates where ITS amplification was unsuccessful, the large subunit (LSU) rRNA region was amplified using primers NL-1 (5'-GCATATCAATAAGCGGAGGAAAAG-3') and NL-4 (5'-GGTCCGTGTTTCAAGACGG-3'). PCR cycling conditions consisted of initial denaturation at 98 °C for 30 s, followed by 35 cycles of 98 °C for 10 s, 60 °C for 30 s, and 72 °C for 1 min, with a final extension at 72 °C for 5 min. PCR products were visualized by agarose gel electrophoresis, and products displaying single, distinct bands were selected for sequencing.

**Sequencing and taxonomic identification.** PCR products (20 µL) were submitted to Eurofins Genomics for purification and Sanger sequencing using primers ITS4 and NL-4 for the ITS and LSU regions, respectively. Resulting sequences were obtained in FASTA format and analyzed using BLAST against the NCBI ITS, LSU, and standard nucleotide databases. For each sequence, the top 3–5 matches from the respective databases were compared based on bit score, query coverage, E-value, and sequence identity. Where database annotations were uninformative (named such as *Ascomycota* sp., *Basidiomycota* sp., or uncultured fungus), additional high-scoring matches were examined to determine the most likely taxonomic identification. Species assignments were guided by a minimum sequence identity threshold of 98.41% to reference sequences. For isolates falling below this threshold, the species name of the closest matching reference sequence was provisionally assigned, and sequence identity values were retained to facilitate future taxonomic investigation.

**Strain cataloguing and representative panel selection.** A database was maintained containing strain identifiers, sample origins, morphological observations, and sequencing results. From the complete collection, 89 strains were selected for phenotypic characterization, typically representing one isolate per identified species, with consideration of taxonomic diversity, morphology, and geographic distribution.

### Growth characterization in microcultures

The 89 representative yeast strains were recovered from cryopreserved stocks by inoculation into 3 mL YPDF medium and incubated at approximately 25 °C with shaking at 150 rpm for 4 days. Cultures were centrifuged at 5,000 × g for 5 min, and the resulting pellets were washed with sterile YP medium (1% yeast extract, 2% peptone) without added sugar. Washed cells were diluted tenfold in YP medium.

Aliquots (10 µL) of the diluted cultures were inoculated into 96-well Growth Profiler microtiter plates (Enzyscreen, catalogue no. CR1496dg) containing 240 µL YP medium supplemented with either 2% fructose or glucose at concentrations of 2%, 10%, 20%, 40%, 60%, 70%, and 80% (w/v). For sugar utilization experiments, strains were additionally cultivated in YP medium containing either 2% glucose + 2% fructose or 10% glucose + 10% fructose (w/v). These cultures were subsequently sampled for HPLC analysis. Plates were incubated in a Growth Profiler 960 (Enzyscreen) at 30 °C with shaking at 250 rpm for 7 days.

Growth was monitored by automated imaging of the microtiter plate wells. Green Values, derived from image analysis, were used as a proxy for biomass accumulation. Growth curves were analyzed to extract parameters including lag phase duration, maximum growth rate, final yield, and area under the curve (AUC) allowing comparison of growth performance across sugar concentrations.

### Honey fermentation assays

Commercial honey (Svensk Honung, Svensk Honungsförädling AB, Sweden) was heat-treated at 90 °C for 1 h to melt the crystalline honey and inactivate endogenous microorganisms, then cooled overnight at room temperature. The honey was diluted to 60%, 70%, 80%, and 90% (v/v) with sterile deionized water and thoroughly homogenized. Aliquots (10 mL) were dispensed into sterile 15 mL tubes.

Ten yeast strains selected for their high osmotolerance were recovered from YPDF agar plates and inoculated in triplicate into each honey dilution using one loopful of cells (approximately 10 µL). Cultures were mixed to distribute the inoculum evenly. Three controls were included: (i) uninoculated heat-treated honey (negative control), (ii) a time-zero (T0) sample, and (iii) actively fermenting honey (positive control). All tubes were sealed and incubated in darkness at 25 °C without agitation. Fermentation was assessed qualitatively by visual inspection for gas formation over 25 days.

In a separate fermentation experiment, *Metschnikowia anglica*, *Zygosaccharomyces siamensis*, and *Entelexis paramagnoliae* were cultivated in 70% honey under the same conditions. Samples were collected on day 21 for chemical analyses. For volatile compound profiling, 3 mL of fermented honey was transferred to 20 mL GC–MS headspace vials. For extracellular metabolite profiling, 5 mL of the corresponding cultures was collected for extraction and UPLC–MS/MS analysis.

### Untargeted LC–MS/MS analysis of fermented honey

Fermented honey samples (5 mL), collected on day 21, were centrifuged at 1,500 rpm for 10 min to remove cells. Supernatants were acidified to pH 3 using glacial acetic acid (99%) and extracted twice with 10 mL ethyl acetate. The organic fractions were evaporated to dryness using a MiVac concentrator and reconstituted in methanol. Metabolite profiling was performed using an ACQUITY Premier UPLC system coupled to a Xevo MRT mass spectrometer (Waters, Milford, MA, USA). Chromatographic separation was achieved using an ACQUITY HSS T3 column (2.1 × 100 mm, 1.8 µm) maintained at 45 °C. Mobile phases consisted of water containing 0.1% formic acid (A) and acetonitrile containing 0.1% formic acid (B), delivered at 0.4 mL/min. The gradient increased from 5% to 100% B over 6 min, remained at 100% B until 10.5 min, and was subsequently returned to initial conditions, giving a total run time of 12 min. Mass spectra were acquired using electrospray ionization in positive and negative modes in separate runs. Data were collected in MSE mode over an m/z range of 50–1,200, using a low-energy function of 6 eV and a high-energy collision-energy ramp of 20–50 eV. Mass accuracy was maintained using lock-mass correction. Raw data were processed using MS-DIAL v4.6 for peak detection, MS/MS deconvolution, retention-time alignment, and export of aligned feature tables and corresponding MS/MS spectra in GNPS format. Downstream processing followed the pipeline developed by D. Brunnsåker (https://github.com/DanielBrunnsaker/bio-project-processing), including blank- and detection-based filtering, QC-based drift correction, probabilistic quotient normalization, correlation-based collapsing of adducts and in-source fragments, spectral library matching, and integration of SIRIUS/CANOPUS predictions. Features were required to have an MS/MS spectrum, a maximum intensity exceeding 10,000, a sample-to-blank ratio greater than 5, and detection in ≥20% of samples and ≥60% of the eight pooled QC injections. Adducts and in-source fragments were collapsed based on correlation (r > 0.9) within a 1-s retention-time window. Of the 3,589 features meeting these criteria, 1,571 with a QC relative standard deviation ≤0.4 and D-ratio ≤0.6 were retained for downstream analyses. Compound classes were predicted using SIRIUS 5.8.6/CANOPUS, retaining NPClassifier pathway assignments with probabilities >0.5.

### GC–MS analysis of volatile compounds

Sterile 20 mL headspace glass vials fitted with sampling screw caps were filled with 3 mL fermented honey collected on day 21. Samples were equilibrated at room temperature for 24 h before analysis. Before extraction, vials were equilibrated at 40 °C for 10 min. Volatile compounds were extracted from the headspace using a polydimethylsiloxane/divinylbenzene/Carboxen (PDMS/DVB/CAR) SPME fiber for 20 min at 40 °C. The fiber was subsequently thermally desorbed in the GC injection port for 5 min at 220 °C in splitless mode. Chromatographic separation was performed using an Agilent GC system equipped with a DB-WAX column (30 m × 0.32 mm × 1 μm), with helium as the carrier gas. The oven temperature was programmed from 35 °C to 280 °C using a multistep gradient. Mass spectrometric detection was performed using a single-quadrupole GC–MS system operated in full-scan mode (m/z 30–300). Compounds were identified based on mass spectra and retention times. Quantification was performed using flame ionization detection (FID).

Apparent odor activity values (OAVs) were calculated as OAV = C/T, where C represents the estimated headspace concentration and T the odor detection threshold in air. Headspace concentrations were estimated by dividing the amount of each compound quantified by FID (ng) by the headspace volume (17 mL), calculated as the difference between vial volume (20 mL) and sample volume (3 mL). Concentrations were expressed as ng/mL, numerically equivalent to $mg/m^3$. Because SPME extracts only a fraction of the available volatile compounds, these values were treated as apparent concentrations, and the resulting OAVs were interpreted as relative estimates rather than absolute odor activity. Odor detection thresholds were obtained from van Gemert [1], using values reported by Nagata [2] where available and otherwise the median of the listed thresholds. For co-eluting 2- and 3-methyl-1-butanol, the threshold of 3-methyl-1-butanol was used. For trimethylbenzene, where the isomer could not be determined, the threshold of 1,2,4-trimethylbenzene was applied. No OAVs were calculated for benzyl alcohol, 3-methyl-1-pentanol, or 4-methyl-1-hexanol because suitable air detection thresholds were unavailable. OAVs were reported as $\log_{10}$(OAV), with values greater than zero considered indicative of potentially odor-active compounds (Table S2).

### Data analysis and visualization

Data processing, statistical analyses, and visualizations were primarily performed in R (version 4.6.1). Data manipulation and visualization used packages from the tidyverse, including *dplyr* and *ggplot2*. Treemaps were generated using *treemapify*, geographic maps using *sf*, *rnaturalearth*, *rnaturalearthdata*, *ggspatial*, and *lwgeom*, and heatmaps using *ComplexHeatmap*. Taxonomic relationships and relative abundances were visualized using *metacoder*. Additional packages included *vegan* for community similarity analysis, *growthcurver* for growth-curve fitting, and *fmsb* for radar plots.

**Community composition and growth analysis.** Species presence–absence matrices were constructed for individual honey batches, and pairwise Jaccard distances were calculated using the *vegan* package. Samples were hierarchically clustered using average linkage, and the resulting similarity matrix was visualized using *ComplexHeatmap*. Growth curves were smoothed using locally estimated scatterplot smoothing (LOESS; span = 0.2) and transformed to monotonically increasing values before fitting logistic growth models using *growthcurver*. Curves with flat lines were excluded from parameter extraction. Growth performance was summarized using the area under the fitted growth curve (AUC), and replicate measurements were averaged for each strain and condition. Relationships between osmotolerance and isolate abundance were examined using linear regression, with AUC at 70% glucose as the explanatory variable and $\log_{10}$-transformed isolate abundance as the response variable.

**Metabolite and volatile compound analysis.** Glucose and fructose consumption were calculated as percentage reductions relative to measured concentrations in uninoculated medium. For volatile compound visualization, abundances were $\log_{10}(x + 1)$-transformed and standardized to row-wise Z-scores before hierarchical clustering. Apparent odor activity values were summed within aroma categories and $\log_{10}$-transformed for radar plot visualization. For LC–MS/MS data, curated feature intensities were $\log_2$-transformed, mean-centered, and Pareto-scaled before principal component analysis (PCA). Missing or non-positive intensities were replaced with half the minimum positive intensity for the corresponding feature. Differences among fermentation treatments were evaluated using permutation-based multivariate analysis of variance (PERMANOVA) based on Euclidean distances, with 9,999 permutations. Homogeneity of multivariate dispersion was also assessed using a permutation-based test.

Illustrative figures were hand-drawn using a Wacom Cintiq 22HD graphics tablet and Clip Studio Paint EX (version 2.0) with custom brushes. Final figure layouts and graphical adjustments were performed using Adobe Illustrator (Creative Cloud 2026).

**Litterature**

1. van Gemert LJ. Odour thresholds : compilations of odour threshold values in air, water and other media. Oliemans Punter, 2011.
2. Nagata Y. Measurement of Odor Threshold by Triangle Odor Bag Method. Odor Measurement Review. Tokyo: Office of Odor, Noise and Vibration, Environmental Management Bureau, Ministry of the Environment, Government of Japan, 2003, 118–127.
