## Supplementary tables for "Nationwide citizen science reveals core and rare osmotolerant yeast communities in Swedish honey"

**Table S1. Species identified from Swedish honey.** The best match to known species were used for species identification.

| Species name | Isolates (n) | Unique sites(n) | Mean % identity | Mean% ITS identity | Mean % LSU identity |
| --- | --- | --- | --- | --- | --- |
| Entelexis paramagnoliae | 881 | 82 | 97.27 | 97.26 | 99.71 |
| Rhodotorula mucilaginosa | 192 | 39 | 99.53 | 99.53 | 99.56 |
| Zygosaccharomyces rouxii | 186 | 39 | 98.36 | 98.05 | 99.77 |
| Starmerella bombi | 142 | 43 | 99.21 | 99.19 | 99.79 |
| Zygosaccharomyces siamensis | 126 | 8 | 99.27 | 98.2 | 99.53 |
| Aureobasidium pullulans | 119 | 25 | 99.31 | 99.35 | 99.23 |
| Metschnikowia anglica | 86 | 11 | 97.89 | 98.26 | 97.25 |
| Filobasidium wieringae | 76 | 20 | 99.35 | 99.27 | 99.61 |
| Starmerella bombicola | 69 | 11 | 99.45 | 99.43 | 99.86 |
| Zygosaccharomyces mellis | 56 | 12 | 99.42 | 98.95 | 99.69 |
| Entelexis sorbosivorans | 48 | 22 | 98.57 | 91.63 | 98.87 |
| Zygosaccharomyces siamensis/mellis | 41 | 1 | 98.58 | 98.58 |  |
| Metschnikowia pulcherrima | 35 | 8 | 97.24 | 96.88 | 98.46 |
| Naganishia albida | 35 | 4 | 99.39 | 99.39 | 99.38 |
| Papiliotrema flavescens | 32 | 3 | 99.26 | 99.22 | 99.91 |
| Aureobasidium pullulans/lini | 24 | 8 | 99.52 | 99.52 |  |
| Entelexis apis | 23 | 8 | 97.19 | 97.41 | 92.56 |
| Zygosaccharomyces pseudorouxii | 23 | 13 | 99.17 | 99.17 |  |
| Metschnikowia sp | 15 | 4 | 98.23 | 98.23 |  |
| Wickerhamomyces anomalus | 14 | 2 | 99.58 | 99.58 |  |
| Sydowia polyspora | 13 | 2 | 99.46 | 99.46 |  |
| Aureobasidium melanogenum | 10 | 7 | 97.67 | 97.67 |  |
| Aureobasidium intercalariosporum | 9 | 7 | 98.2 | 98.2 |  |
| Debaryomyces sp | 9 | 3 | 99.56 | 99.56 |  |
| Metschnikowia colchici | 9 | 1 | 96 | 95.73 | 98.16 |
| Candida parapsilosis | 8 | 4 | 99.27 | 99.19 | 99.82 |
| Naganishia diffluens/albida | 8 | 1 | 99.47 | 99.47 |  |
| Naganishia globosa | 8 | 5 | 99.72 | 99.72 |  |
| Rhodotorula dairenensis | 8 | 5 | 99.66 | 99.66 | 99.64 |
| Torulaspora delbrueckii | 8 | 2 | 99.64 | 99.64 |  |
| Debaryomyces hansenii | 6 | 3 | 99.66 | 99.66 |  |
| Neodothiora populina | 6 | 4 | 98.52 | 98.51 | 98.57 |
| Coniochaeta hoffmannii | 5 | 1 | 99.28 | 99.28 |  |
| Naganishia antarctica | 5 | 3 | 99.02 | 99.02 |  |
| Naganishia sp | 5 | 5 | 99.24 | 99.24 |  |
| Coniochaeta ligniaria | 4 | 1 | 98.31 |  | 98.31 |
| Cystobasidium laryngis | 4 | 2 | 99.86 | 99.86 |  |
| Naganishia friedmannii | 4 | 4 | 99.56 | 99.38 | 99.74 |
| Rhodotorula sp | 4 | 3 | 98.64 | 98.64 |  |
| Aureobasidium sp | 3 | 3 | 97.31 | 93.38 | 99.27 |
| Coniochaeta nivea | 3 | 1 | 99.38 | 99.38 |  |
| Dothiora phillyreae | 3 | 2 | 97.67 | 97.67 |  |
| Hormonema macrosporum/Sydowia polyspora | 3 | 2 | 99.58 |  | 99.58 |
| Naganishia diffluens | 3 | 3 | 99.25 | 99.25 |  |
| Rhodotorula graminis | 3 | 2 | 99.7 | 99.64 | 99.82 |
| Starmerella sp | 3 | 3 | 96.66 | 96.66 |  |
| Coniochaeta cymbiformispora | 2 | 1 | 97.63 | 97.63 |  |
| Coniochaeta prunicola | 2 | 1 | 98.44 |  | 98.44 |
| Cystobasidium slooffiae | 2 | 1 | 99.63 | 99.63 |  |
| Exophiala xenobiotica | 2 | 2 | 99.83 | 99.83 |  |
| Metschnikowia chrysoperlae | 2 | 1 | 98.47 | 97.55 | 99.38 |
| Metschnikowia rubicola | 2 | 1 | 97.58 |  | 97.58 |
| Meyerozyma caribbica | 2 | 1 | 99.82 |  | 99.82 |
| Meyerozyma guilliermondii | 2 | 2 | 97.84 | 97.84 |  |
| Aureobasidium insectorum | 1 | 1 | 99.25 | 99.25 |  |
| Aureobasidium leucospermi/subglaciale | 1 | 1 | 99.46 |  | 99.46 |
| Aureobasidium namibiae | 1 | 1 | 98.84 | 98.84 |  |
| Blastobotrys arbuscula | 1 | 1 | 99.2 | 99.2 |  |
| Candida argentea | 1 | 1 | 98.93 | 98.93 |  |
| Coniochaeta africana | 1 | 1 | 97.99 |  | 97.99 |
| Curvibasidium cygneicollum | 1 | 1 | 99.11 | 99.11 |  |
| Cystobasidium benthicum | 1 | 1 | 98.11 |  | 98.11 |
| Cystobasidium ritchiei | 1 | 1 | 95.54 | 95.54 |  |
| Dothiora europaea | 1 | 1 | 99.29 |  | 99.29 |
| Exophiala dermatitidis | 1 | 1 | 99.29 |  | 99.29 |
| Filobasidium chernovii | 1 | 1 | 95.08 | 95.08 |  |
| Filobasidium oeirense | 1 | 1 | 100 | 100 |  |
| Filobasidium uniguttulatum | 1 | 1 | 98.79 | 98.79 |  |
| Metschnikowia pulcherrima/chrysoperlae | 1 | 1 | 97.51 | 97.51 |  |
| Metschnikowia ziziphicola | 1 | 1 | 94.64 | 94.64 |  |
| Metschnikowia ziziphicola/pulcherrima | 1 | 1 | 97.54 | 97.54 |  |
| Naganishia adeliensis | 1 | 1 | 99.65 | 99.65 |  |
| Naganishia bhutanensis | 1 | 1 | 99.12 | 99.12 |  |
| Naganishia liquefaciens | 1 | 1 | 99.66 | 99.66 |  |
| Naganishia uzbekistanensis | 1 | 1 | 99.14 | 99.14 |  |
| Neodothiora pruni | 1 | 1 | 98.65 | 98.65 |  |
| Papiliotrema laurentii | 1 | 1 | 99.83 |  | 99.83 |
| Papiliotrema terrestris | 1 | 1 | 97.26 |  | 97.26 |
| Piskurozyma capsuligena | 1 | 1 | 94.33 | 94.33 |  |
| Rhodosporidiobolus colostri | 1 | 1 | 100 | 100 |  |
| Rhodotorula babjevae | 1 | 1 | 99.1 | 99.1 |  |
| Rhodotorula graminis/glutinis | 1 | 1 | 99.45 | 99.45 |  |
| Scleroconidioma sphagnicola | 1 | 1 | 96.04 | 96.04 |  |
| Starmerella apicola | 1 | 1 | 100 |  | 100 |
| Ustilentyloma graminis | 1 | 1 | 99.65 | 99.65 |  |

**Table S2.** Log₁₀-transformed odor activity values (log₁₀OAV) of volatile compounds detected in the headspace of the honey samples. OAV = concentration / odor threshold, with concentration and threshold both expressed in mg m⁻³; log₁₀ OAV = 0 marks the detection threshold.

|  |  |  | **log₁₀(OAV)** | | | |
| --- | --- | --- | --- | --- | --- | --- |
| **Compound** | **Aroma descriptor** | **Odor threshold (mg m⁻³)** | **Control** | ***M. anglica*** | ***E. paramagnoliae*** | ***Z. siamensis*** |
| ***Alcohols*** |  |  |  |  |  |  |
| 1-Propanol | Alcoholic | 0.24 | – | – | 1.43 | – |
| 2- and 3-Methyl-1-butanol^c^ | Alcoholic | 0.0061 | 2.24 | 3.49 | 4.70 | 2.84 |
| 2-Methyl-1-propanol | Alcoholic | 0.033 | – | 1.86 | 3.40 | – |
| 2-Phenylethanol^a^ | Floral | 0.031 | 1.77 | 3.13 | 3.79 | 1.99 |
| 3-Methyl-1-pentanol | Alcoholic | n/a | – | n/a | n/a | n/a |
| 4-Methyl-1-hexanol | Sweet | n/a | – | – | n/a | – |
| Benzyl alcohol | Floral | n/a | n/a | n/a | n/a | n/a |
| ***Aldehydes*** |  |  |  |  |  |  |
| Benzaldehyde^a^ | Almond | 1.6 | 0.48 | – | – | *−0.07* |
| Benzeneacetaldehyde^a^ | Almond | 0.00072 | 3.82 | 3.86 | – | 3.75 |
| Furfural^a^ | Almond | 0.84 | 0.26 | – | – | – |
| Nonanal | Floral | 0.002 | 3.02 | – | – | – |
| ***Acids*** |  |  |  |  |  |  |
| 2-Methylpropanoic acid | Acetic | 0.0054 | – | 2.54 | – | – |
| ***Esters*** |  |  |  |  |  |  |
| Ethyl acetate | Fruity | 3.1 | – | – | 0.02 | – |
| ***Ketones*** |  |  |  |  |  |  |
| 4-Methyl-2-pentanone | Fruity | 0.7 | – | – | – | 0.15 |
| ***Sulfur compounds*** |  |  |  |  |  |  |
| Dimethyl disulfide | Sweet | 0.0084 | 1.98 | – | 1.64 | – |
| Dimethyl trisulfide^b^ | Sweet | 0.0075 | 1.87 | – | – | – |
| ***Hydrocarbons*** |  |  |  |  |  |  |
| Octane | Fuel-like | 8 | *−0.19* | – | – | – |
| Styrene | Sweet | 0.15 | 0.57 | – | – | – |
| Trimethylbenzene | Fuel-like | 0.59 | *−0.03* | 0.01 | 0.39 | *−0.05* |
| ***Terpenoids*** |  |  |  |  |  |  |
| Linalool^a^ | Floral | 0.063 | 1.34 | – | 1.37 | – |

Thresholds are air detection thresholds from Nagata (2003), as compiled by van Gemert (2011), unless marked. –, not detected in the sample; n/a, no air threshold available, so OAV could not be calculated. Values in italics are below 0, i.e. below the detection threshold (OAV below 1).

^a^ Median of the detection thresholds listed in van Gemert (2011) for air (benzaldehyde, 6 values; benzeneacetaldehyde, 1 value; furfural, 2 values; linalool, 3 values; 2-phenylethanol, 8 values).

^b^ No labelled detection threshold available; median of three unlabelled values in van Gemert (2011). Low confidence.

^c^ Co-eluting 2- and 3-methyl-1-butanol; threshold of the 3-methyl isomer.
